# Enhanced Detection of Age-related Macular Degeneration in Low-quality Retinal Images via Noise-Augmented YOLO and Adaptive Attention Mechanisms

**DOI:** 10.64898/2026.08.04.742925

**Authors:** Xidong Bai, Kazumasa Kishimoto, Osamu Sugiyama, Hiroshi Tamura

## Abstract

This study aims to improve the detection performance of age-related macular degeneration (AMD) in low-quality retinal images.

**Background:** AMD is a leading cause of vision loss among older adults globally, and accurate detection is crucial for clinical management. However, low-quality optical coherence tomography (OCT) images significantly compromise diagnostic accuracy.

**Objective:** To enhance AMD detection in low-quality images using noise-augmented data augmentation and an improved YOLO deep learning model.

**Methods:** Public datasets from UCSD and Duke University were utilized; the training dataset comprised 24,980 OCT images (high-quality and noise-augmented low-quality), while the testing dataset included 1,000 images (584 AMD, 416 normal). The model is based on the YOLOv8n framework, integrated with Squeeze-and-Excitation blocks (SEblock) and Adaptive Sparse Self-Attention (ASSA), with an additional 160×160 detection layer for detecting small lesions. Evaluation metrics included accuracy, sensitivity, specificity, and F2-score.

**Results:** The proposed model achieved an accuracy of 99.02%, sensitivity of 98.17%, specificity of 100%, and an F2-score of 98.50% on the Duke dataset. Detection rates were significantly improved compared to traditional methods, particularly in low-quality images, with a detection rate of 89.60%, markedly superior to original YOLOv8n (55.10%) and classical models like ResNet50.

**Conclusion:** The enhanced model, employing noise-augmented training data and improved attention mechanisms, demonstrates excellent AMD detection capabilities in low-quality OCT images, showing broad potential for clinical applications.

## 1. Introduction

Age-related macular degeneration (AMD) is a leading cause of vision loss in elder people across all income levels and constitutes 6% to 9% of global legal blindness. The number of affected individuals is projected to rise from 196 million in 2020 to 288 million in 2040 worldwide[1]. Optical coherence tomography has become the most common noninvasive imaging modality for detecting AMD. Early and accurate detection is crucial for effective management and treatment. However, current diagnoses rely on high-quality OCT images, which may not always be available. Especially in patients with ocular comorbidities such as cataracts, even state-of-the-art OCT systems may not generate clear images. Given the accelerating trend of population aging, it is anticipated that the need to make clinical judgments based on noisy images will persist for the foreseeable future. OCT images of low quality often exhibit lower contrast, brightness, and more noise, which significantly impact diagnostic accuracy and reliability. Thus, enhancing AMD detection performance on low-quality retinal images remains a challenge in clinical ophthalmology.

Traditionally, convolutional neural network (CNN) based detection methods also yielded encouraging results. Lee et al. used a modified Visual Geometry Group (VGG) model to classify OCT Images of AMD patients and normal people[2]. Ruchir Srivastava et al. used the Resnet50 model and preprocessed images to identify AMD patients and the normal, obtaining advanced performance[3]. There are other works related to automatic detection based on OCT Images. Karri et al. used the GoogleNet model with transfer learning to identify AMD patients and DME (diabetic macular edema) patients[4]. Alqudah et al. proposed AOCT-Net for multiclass classification (CNV, DME, AMD)[5].

You Only Look Once(YOLO) is known for its swift and high accuracy. However, the detection rate of the original version of YOLOv8n degraded when applied to low-quality OCT images, primarily due to their insufficient handling of noise and poor-quality data.

In this study, we propose a novel redesigned detection model based on YOLOv8n, integrating with Squeeze-and-Excitation network(SEblock)[6]and Adaptive Sparse Self-Attention (ASSA) modules[7], which show significant improvements in contextual feature learning and noise reduction. These methods selectively emphasize useful feature channels and minimize redundant information. Moreover, we performed noise-added image augmentation on the high-quality images in the training datasets to minimize the gap between high-quality OCT images and low-quality OCT images.

## 2. Method

In order to simulate a situation in which the model trained on high-quality images recognizes images of low quality. We utilize a refined YOLOv8 model with Squeeze-and-Excitation networks and Adaptive Sparse Self-Attention, trained on OCT images from the University of California, San Diego (UCSD).

**Fig. 1.**
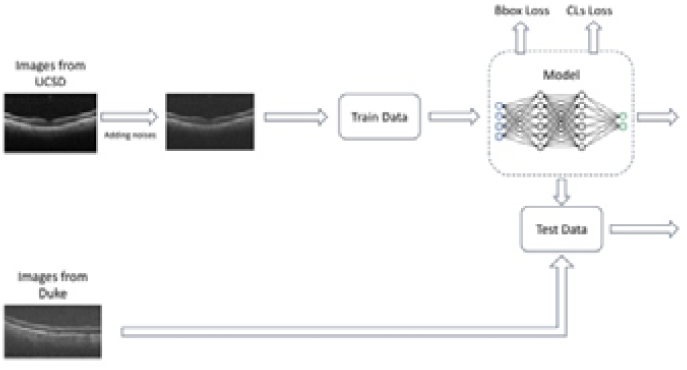
Process of the automatic detection model.

### 2.1 Datasets

In the current study, we used open datasets from Duke University and UCSD. Datasets from Duke University[8] contain 1000 images (584 AMD images and 416 Normal images) for testing. Datasets from UCSD[9] contain 24980 images(6265 AMD,16865 Normal images,350 Normal images with noise, and 1500 AMD images with noise) for training.

**Fig. 2.**
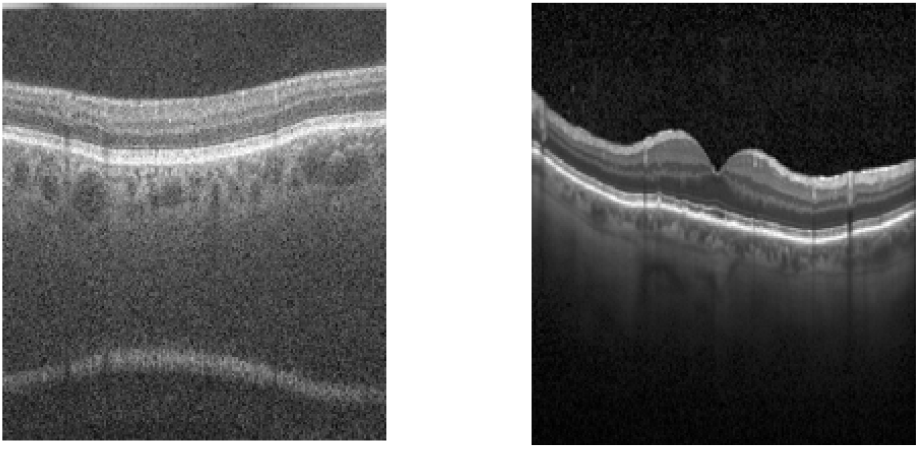
(a) An image from the dataset of Duke University (b) An image from the dataset of UCSD.

For better simulation, we add noise to the original images to augment the low-quality image of the training data. We add Gaussian noise to the image, reducing the contrast as well as increasing the brightness.

Our model was rigorously tested on open datasets from Duke University and the University of California, San Diego, demonstrating superior performance compared to traditional approaches and highlighting its potential for improving AMD detection in real-world clinical practice.

### 2.2 YOLO

YOLO is a deep-learning algorithm, with the first version released in 2015 and developing 12 versions so far, which can simultaneously localize and classify targets. The core concept of YOLO is using a convolutional neural network (CNN) to predict the category of each target and the location of the bounding box[10].In the current study, we improve the model based on YOLOv8n, and the new model has better performance in detecting OCT images.

### 2.3 Squeeze-and-Excitation Networks (SEbolck)

SEbolck was released in 2017[6], introducing an adaptive channel attention mechanism for improving the performance of the model by focusing on more crucial feature channels. The core steps of SEblock include Squeeze and Excitation.

**The Squeeze part** squeezes the feature maps into a channel vector by Global Average Pooling. H and W remain spatial dimensions, and c is the number of channels.

**The excitation part** generates the weights of each channel by a fully connected layer and an activation function(ReLU). The weights are compressed to the range[0,1] by a Sigmoid function. The process of Excitation can be represented by the equation:

Finally, the **recalibration** is achieved by multiplying the weights with the input feature maps.

Compared to other attention mechanisms, SEblock has less computational overhead and can be easier to be integrated into other models, which could significantly improve the performance of the model[6].

### 2.4 Adaptive Sparse Self-Attention(ASSA)

ASSA is a novel approach to image restoration based on the Transformer architecture. The key target is enhancing the ability to recover clear images from degraded images by an efficient self-attention mechanism and reducing redundancy in feature maps. The ASSA includes two branches: **Sparse Self-Attention (SSA)**and **Dense Self-Attention(DSA)**.

The equations of Query, Key, and Value of ASSA:

Where X is the input feature map, and W_Q, W_K, W_V are the linear projection matrices for queries, keys, and values.

**Sparse Self-Attention** refers to sparsification in the self-attention mechanism by only focusing on the interactions between the most relevant tokens. This mechanism reduces computational resources and avoids introducing redundant information. Sparse self-attention uses ReLU^2^ to filter out irrelevant information by removing low-matched query-key pairs.

**Dense Self-Attention** is the standard self-attention mechanism that each query token has interactions with all key tokens, which may introduce redundant information.

Where d is the dimension of queries and keys, and B is the learnable relative positional bias.

**ASSA** combines the sparse and dense branches adaptively by introducing 2 learnable weights w1 and w2. Where V is the value matrix.

The weights w1 and w2 are computed by the following formula:

Where a1 and a2 are learnable parameters related to the sparse and dense attention. N is the total number of branches[7].

### 2.5 Optimizer

To keep better performance, we use Stochastic Gradient Descent(SGD) instead of adaptive optimization. Although adaptive methods converge faster, SGD and SGD with momentum perform better than adaptive optimization in all models and tasks[11].

### 2.6 Detection Rate

For the initial version of YOLOv8n, only a few images from the Duke datasets can be recognized and then classified because of their low quality. In this paper, we define detection rate as the ratio of recognized images to total image samples.

### 2.7 Redesigned detection model

The initial version of YOLOv8 performs well in classifying AMD OCT images. However, the detection rate is only 16%. In order to improve the detection rate, we introduce new SEblock modules and ASSA attention mechanism in YOLOv8n.

For the original YOLOv8n, the neck fuses the extracted features from the backbone part and outputs three scales of feature maps, 20×20, 40×40, and 80×80, which are used for detecting targets with different scales. The 20×20 detection layer is used to recognize large targets with a size of 32×32 or more, while the 40×40 detection layer is designed for detecting medium-sized targets of 16×16, and the 80×80 detection layer is used to detect smaller targets of 8×8. When the Upsampling multiplier of the neck is large, the deeper feature maps will lose detailed information on the small targets, resulting in difficulty in classification for small target samples. For some nuances on the surface of OCT images, the 3 original detection layers cannot successfully recognize them, which may be the reason for missed detection. Thus, we propose adding a 160×160 detection layer based on the initial version to detect tiny targets smaller than 8×8.

As shown in Fig.3, compared with the original version of YOLOv8n, the neck part is added with another Upsampling operation behind the second Upsampling module to obtain a 160×160 feature map, which is spliced with the 160×160 feature map of the backbone part to obtain a new prediction scale. The SEblock is added among the different sizes of the detection layer for better feature extraction. We add the ASSA module to the Neck part to improve the performance of crawling details for low-quality images. A new detection layer is added to the improved model to enhance the recognition of image details.

**Fig. 3.**
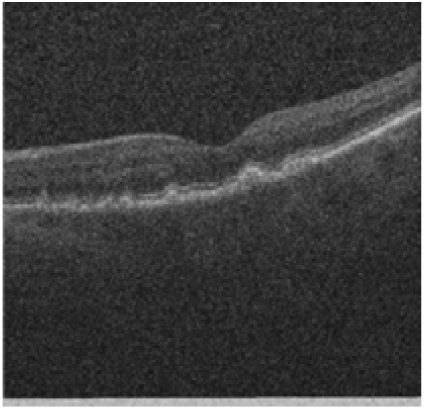
Augmented Images with noise as low-quality images. Augmented Images are based on the University of California, San Diego dataset, with contrast reduced by 30%, brightness increased by 20 units, and Gaussian noise added (standard deviation = 50)

**Fig. 4.**
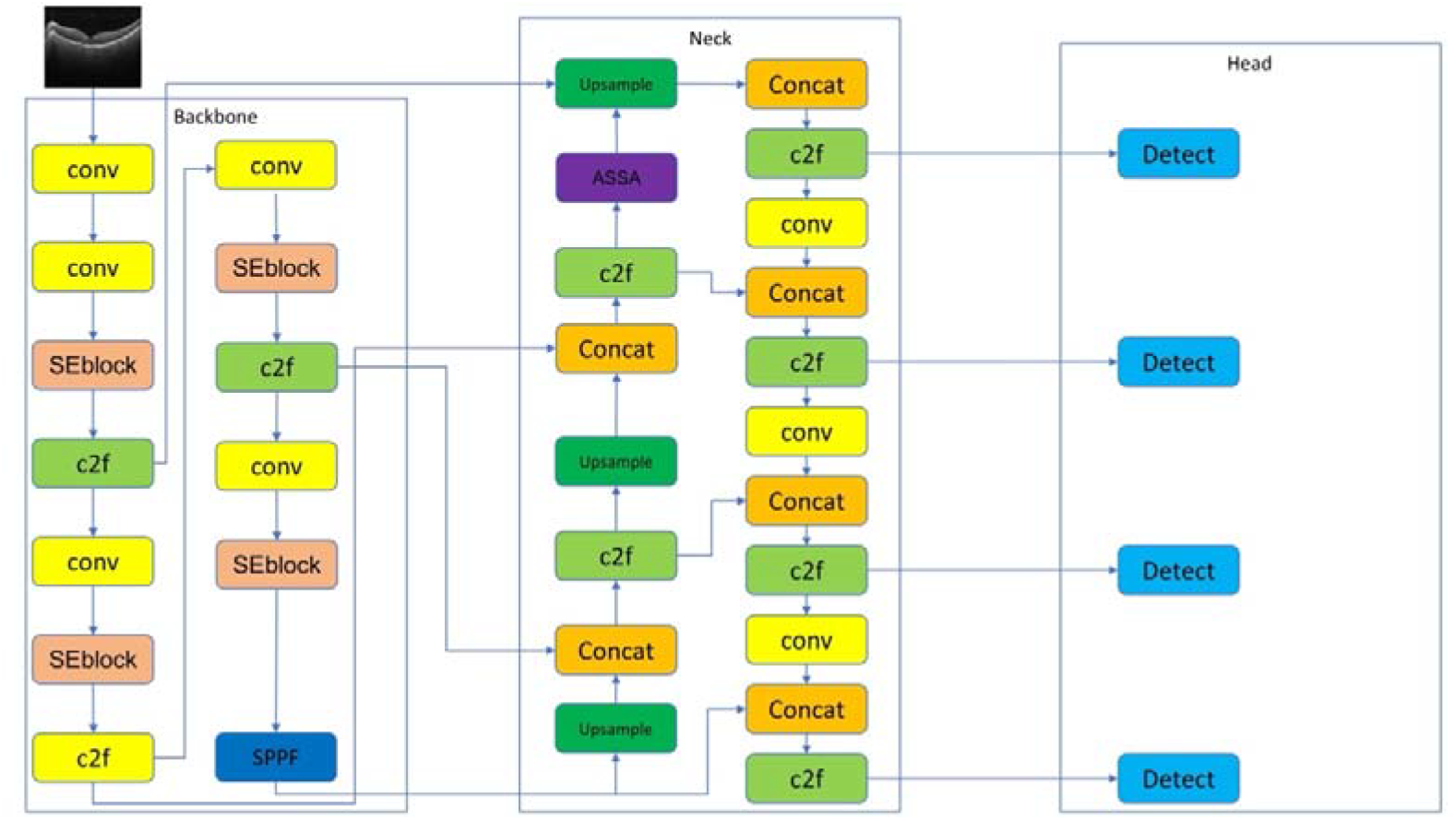
Redesigned Model with SEblock, ASSA module, and additional detect layer.

### 2.8 Ablation Experiments

In order to verify the effect of each module and the differences between datasets on the results. We carried out 2 ablation experiments.

The first experiment is to verify the effect of each module: we set the model into four groups. The first group is the original version of YOLOv8n, the second group is only added SEblock, the third group is not only added the SEblock but also an extra detection layer, the fourth group is based on the second with the addition of ASSA module in the redesigned Neck part.

The second experiment to verify the effect of datasets: we use the UCSD datasets and the UCSD datasets with noise to train the model, respectively, to test the influence of datasets.

### 2.9 Ablation Experiments

The new model showed strong performance and is measured by the accuracy, sensitivity, specificity, and F-2 score compared with the prior studies.

## 3. Results

As shown in Table 1, the proposed model achieved 3.2 percentage points improvement compared with Resnet50, and correctly classified 100% Normal OCT images, which is superior to Resnet at 3.09 percentage points, at the expense of parameters of 11.4 M.

**Table 1.** Comparison with existing models.

| Model | Accuracy | Sensitivity | Specificity | Parameters | F1-score | F2-score |
| --- | --- | --- | --- | --- | --- | --- |
| Modified VGG16[2] | 87.63% | 84.63% | 91.54% | 138M | / | / |
| Resnet50[3] | 95.82% | 95.45% | 96.91% | 25.6M | / | / |
| SE and ASSA added* | 99.02% | 98.17% | 100% | 37M | 99.08% | 98.50% |

\*proposed model in the current study

### 3.1 Results of Ablation Experiments

In the first ablation experiment, we use the same samples, including 416 normal OCT images and 584 AMD images, for testing. These images are all from the Duke datasets. The results are shown in Table 2. According to the results, even the original yolo model shows a relatively good performance in terms of accuracy. Compared with Resnet50, the best model in the prior studies also improves by 4 percentage points.

**Table 2.** Results of the 1st ablation experiment.

| Model | Accuracy | Sensitivity | Specificity | Detection Rate | F2-score |
| --- | --- | --- | --- | --- | --- |
| YOLOv8n | 99.64% | 99.52% | 100% | 55.10% | 99.64% |
| Group2* | 98.72% | 97.72% | 100% | 87.30% | 98.20% |
| Group3* | 98.99% | 98.14% | 100% | 89.10% | 98.47% |
| SE and ASSA added YOLOv8n** | 99.02% | 98.17% | 100% | 89.60% | 98.50% |

\* Group 2 only added SE blocks. Group 3 is not only added SEblock but also an extra detection layer.
\*\*proposed model in the current study
\*\*\*Total number of testing samples is 1000 (416 Normal OCT and 584 AMD OCT).
\*\*\*\*All indicators are based on OCT images that can be recognized.

**Table 3.**
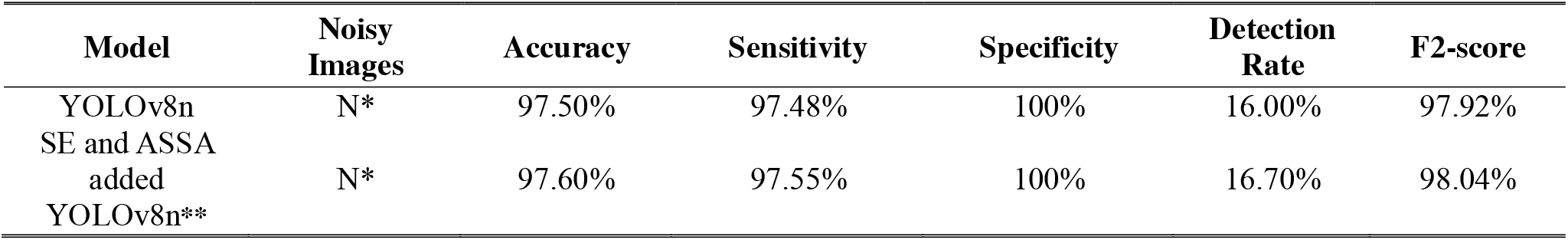
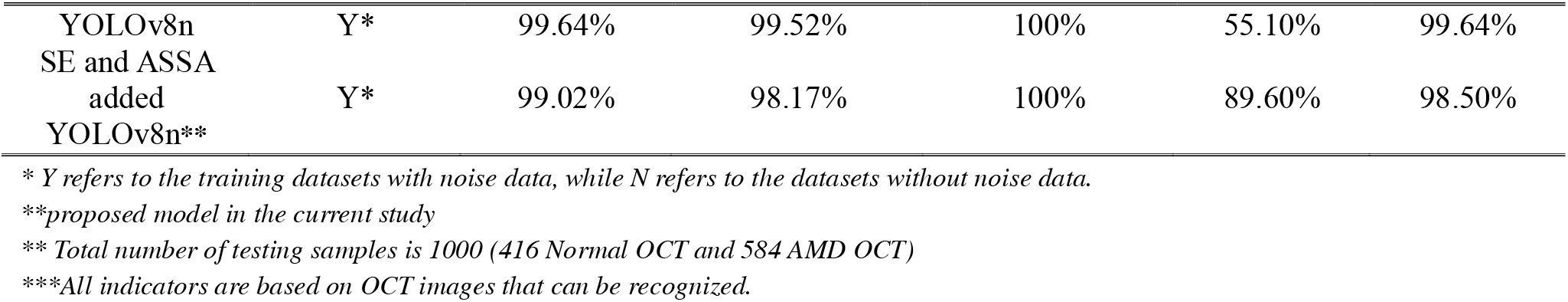
Results of the 2nd ablation experiment.

However, in terms of the detection rate, our model successfully recognizes 896 OCT images(350 Normal images and 546 AMD images). Among the successfully detected OCT images, our model misdiagnosed 10 AMD images, and for normal images, there was no misdiagnosis. The specific performance of the other groups in the first ablation experiment is shown in Table 2.

The results show that our model gains the best performance in all indicators, especially in recognizing AMD images. The original version of YOLOv8n successfully recognized 551 OCT images (AMD: 416, Normal: 135). Group 2 successfully recognized 873 OCT images (AMD: 526, Normal: 347). Group 3 successfully recognized 891 OCT images (AMD: 537, Normal: 354). Our model successfully recognized 896 OCT images (AMD: 584, Normal: 350)

According to the results in the 2^nd^ ablation experiment, the original version of YOLOv8n successfully recognized 160 OCT images(AMD:159, Normal: 1) trained on datasets without noisy data. Our model successfully recognized x OCT images (AMD: 163, Normal: 5).

## 4. Discussion

In this study, we propose a modified YOLOv8n detection model combining Squeeze-and-Excitation (SE) blocks[6], an Adaptive Sparse Self-Attention (ASSA) module[7], and an additional 160×160 detection layer, aiming to improve the ability to recognize age-related macular degeneration (AMD) from low-quality OCT images. The results demonstrated the effectiveness of our design, particularly in increasing detection robustness, accuracy, and sensitivity under degraded imaging conditions.

Our model achieves a detection rate of 99.08%, an accuracy of 99.02% and an F2-score of -score of 98.50%, outperforming classical deep learning models from prior studies (e.g., Resnet50[3], VGG16[2]) and the original version of YOLOv8n. Through ablation studies, we verified the individual effect of each module. The inclusion of the SEblock, the ASSA module in the redesigned neck architecture, and the additional detection layer improved the recognition of AMD lesions and achieved better performance in all indicators.

The results demonstrate that the detection rates of the two models in the training datasets without noisy data are at a low level, while the detection rates of the two models in the training set with noisy data increase significantly, and our model outperformed the original version of YOLOv8n. Moreover, the two models trained without noisy images are not sensitive to normal images. Due to the relatively small sample size of successful detections, we do not consider the indicators of the two models to be trustworthy in the absence of noisy data in the training datasets.

Compared to prior studies, our approach addressed a core problem: the models he model’s performance on low-quality image data. Classic models degrade when applied to low-quality images. By introducing image augmentation (synthetic Gaussian noise, brightness, and contrast) to simulate historical low-quality data, we narrowed the domain gap between training and deployment environments, thus improving the detection rate on low-quality images.

In the mechanism, SEblock emphasizes the most important channels without increasing large number of computational cost. The ASSA module, integrating sparse and dense self-attention, reduced redundancy and preserved relevant local structures. An addition of the 160×160 detection head further enabled the model to capture minute features, which are often critical for early AMD diagnosis.

The F1-score is commonly used to evaluate the performance of classification models. However, in the field of medicine, models focusing on diseases(e.g., polyp[12] and lung diseases[13]) use F2-score as a reference for measuring, since F2-score gives higher weight to recall, especially when the recall is more important than precision, which makes the model more inclined to reduce the number of missed detections (False Negatives).

Although we achieved relatively promising results, several limitations still remain. First, although we utilized noise augmentation to simulate low-quality images from Duke datasets, we still didn’t simulate all situations encountered in real clinical practice. Second, our results were limited to two datasets(UCSD and Duke), and further external validation is needed to ensure generalizability.

## Conclusion

In this study, we proposed a modified YOLOv8n-based AMD detection model, aiming to enhance the detection rate of Age-related Macular Degeneration (AMD) in low-quality retinal OCT images.

By introducing SEblock, ASSA module, and integrating noise augmentation, our model significantly improved the detection rate of low-quality images while outperforming other prior studies in terms of indicators

In future work, we plan to use randomized Gaussian noise and contrast adjustments to simulate a broader range of conditions and to design new models to further enhance robustness. Indeed, due to the frequent occurrence of cataracts and similar conditions, there will likely remain a need to perform fundus diagnoses using high-noise images—not only for AMD, but also for diabetic retinopathy, retinal vascular occlusion, and other retinal diseases. Our proposed method is expected to contribute meaningfully in these challenging clinical scenarios.

## Ethics declaration

All investigations adhered to the principles of the Declaration of Helsinki. Ethical review was exempt because we used two open datasets: Duke University and UCSD. These datasets are open to the research community and contain no identifiable personal information.

## Acknowledgements

This work did not receive any specific grant from funding agencies in the public, commercial, or not-for-profit sectors.

